# AMPK activation promotes transcriptional activation of TFEB through its dephosphorylation

**DOI:** 10.1101/2024.10.22.619589

**Authors:** Florentina Negoita, Conchita Fraguas Bringas, Kristina Hellberg, Katarzyna M. Luda, Hongling Liu, Joyceline Cuenco, Jin-Feng Zhao, Gajanan Sathe, Gopal Sapkota, Kei Sakamoto

**Author notes:** Contact Kei Sakamoto, Novo Nordisk Foundation Center for Basic Metabolic Research, University of Copenhagen, Blegdamsvej 3B, Copenhagen 2200, Denmark.

## Abstract

Transcription Factor EB (TFEB) is a critical regulator of lysosomal biogenesis, autophagy and energy homeostasis through controlling expression of genes belonging to the coordinated lysosomal expression and regulation network. AMP-activated protein kinase (AMPK) has been reported to phosphorylate TFEB at three conserved C-terminal serine residues (S466, S467, S469) and these phosphorylation events were essential for transcriptional activation of TFEB. In sharp contrast to this proposition, here we demonstrate that AMPK activation leads to dephosphorylation of the C-terminal sites, and that AMPK is dispensable for mTORC1-mediated/torin1-sensitive TFEB activation. We show that a synthetic peptide encompassing C-terminal serine residues of TFEB is a poor substrate of AMPK. Treatment of cells with AMPK activator (MK-8722) or mTOR inhibitor (torin1) robustly dephosphorylated TFEB not only at mTORC1-targeted N-terminal serine sites, but also at the C-terminal sites. Loss of function of AMPK abrogated MK-8722-but not torin1-induced dephosphorylation and induction of the vast majority of TFEB target genes.

## Introduction

Transcription factor EB (TFEB) and related transcription factor binding to IGHM enhancer 3 (TFE3) are members of the basic helix-loop-helix leucine zipper family of transcription factors [1]. TFEB is a master regulator of lysosomal biogenesis, autophagy and cellular energy homeostasis through controlling expression of genes belonging to the coordinated lysosomal expression and regulation (CLEAR) network [2,3]. An important mechanism by which TFEB is regulated involves its shuttling between the surface of lysosomes, the cytoplasm, and the nucleus. Such dynamic changes in subcellular localization occur in response to nutrient fluctuations and various forms of cellular and energetic stresses, and are primarily mediated via changes in the phosphorylation of multiple conserved residues in TFEB [1,4] (**Fig. 1A**). Under nutrient- and energy-repleted states, TFEB is held inactive in the cytosol by mechanistic target of rapamycin complex 1 (mTORC1) through phosphorylation of multiple serine residues (including S122, S142, S211) [1]. Upon inhibition of mTORC1 and/or activation of 5’-adenosine monophosphate-activated protein kinase (AMPK) in response to nutrient or energy depletion, TFEB undergoes dephosphorylation and nuclear translocation [5–7]. In the nucleus, TFEB directly binds to a specific E-box sequence (known as CLEAR motif) in the proximal promoters of numerous genes that regulate lysosomal and metabolic functions [2,3]. Mechanistically, it has been reported that activated AMPK promotes nuclear translocation of TFEB through inhibition of mTORC1 via phosphorylation of Regulatory-associated protein of mTOR (Raptor) and tuberous sclerosis complex 2 (TSC2) [8], or via phosphorylation of Folliculin-interacting protein 1 (FNIP1) and the resulting dissociation of Ras-related GTP-binding protein C (RagC), mTORC1 and TFEB from the lysosome [5].

**Figure 1.**
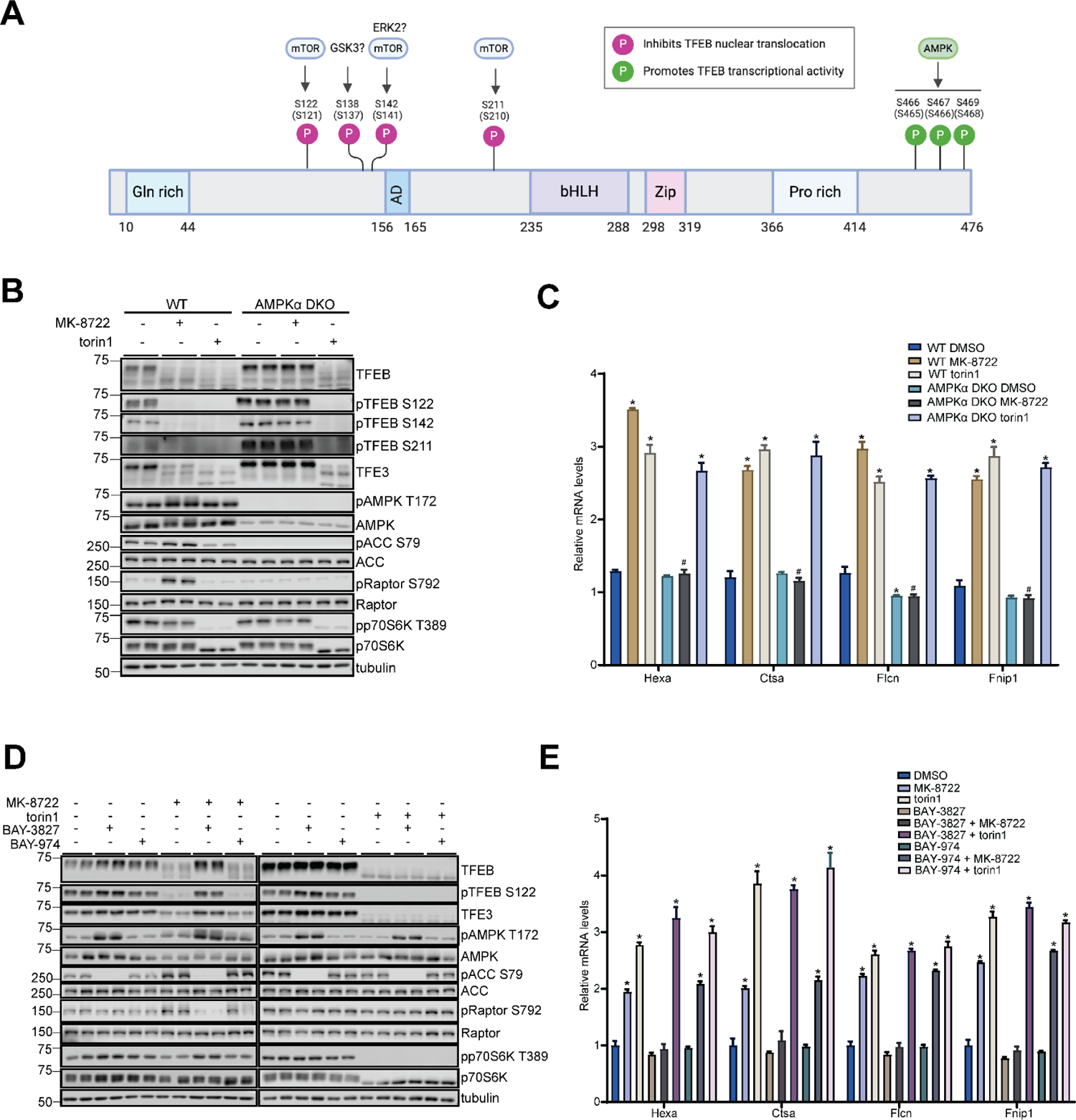
AMPK is dispensable for mTORC1-regulated (torin1-sensitive) transcriptional activity of TFEB. (A) Schematic domain structure and reported phosphorylation sites (validated by phospho-specific antibodies) of human TFEB. Mouse TFEB phosphorylation sites in bracket. Illustration was created with BioRender.com. (B and C) WT and AMPKα1/α2 DKO MEF were treated with vehicle (0.1% DMSO), 10 μM MK-8722 or 100 nM torin1 for 6 h. The extracted proteins and RNA were subjected to immunoblot analysis with the indicated antibodies or RT-qPCR assay, respectively. Representative immunoblot images and qPCR data from three independent experiments are shown. n=2 per treatment condition for immnoblot and n=3 per treatment condition for qPCR. Data in (C) and (E) are expressed mean ± S.E.M. (D and E) WT MEF were pre-incubated with vehicle, 5 μM BAY-3827 or 5 μM BAY-974 for 30 min followed by treatment with 10 μM MK-8722 or 100 nM torin1 for 6 h. Representative immunoblot images and qPCR data from three independent experiments are shown. n=2 per treatment condition, n=3 per treatment condition. Data from one independent experiment is shown. A two-way ANOVA (C) or one-way ANOVA (E) with Šídák’s multiple comparison was performed (* *p* < 0.05 vehicle *vs.* treatment and # *p* < 0.05 KO *vs.* WT).

Notably, AMPK has been reported to phosphorylate TFEB at a cluster of three conserved C-terminal serine residues (S466, S467, S469) (**Fig. 1A**), and these phosphorylation events are required for transcriptional activation of TFEB [7]. The study showed that in AMPK-null cells, while inhibition of mTORC1 by starvation or torin1 (a potent and selective ATP-competitive mTOR inhibitor) promoted TFEB localization to the nucleus, TFEB activity was abrogated. In addition, cellular introduction of a GFP-tagged TFEB mutant which carried mutations in the three C-terminal serine sites to alanine (S466/467/469A) did not disrupt the ability to promote nuclear translocation of TFEB, but abrogated transcriptional activity of TFEB upon activation of AMPK (with 5-aminoimidazole-4-carboxamide ribonucleotide (AICAR)) or inhibition of mTORC1 (with torin1) [7]. However, the mechanism by which AMPK-mediated phosphorylation regulates TFEB activity or whether phosphorylation of all three residues is required is unknown. Given that all three of these tightly clustered C-terminal serine residues of TFEB poorly match the AMPK substrate consensus motif [9,10], it is unlikely that AMPK phosphorylates all three sites (especially with the same stoichiometry). In the current study, we developed a series of phospho-site specific antibodies and synthetic peptides targeting/encompassing the C-terminal serine residues of TFEB and assessed the ability of AMPK to phosphorylate these sites in a cell-free assay, and studied phosphorylation and activity of TFEB in response to a highly-specific allosteric AMPK activator (MK-8722) and torin1 in cells genetically lacking AMPK or treated with a potent and highly-selective AMPK inhibitor (BAY-3827). In stark contrast to the previous study [7], we here demonstrate that AMPK activation promotes *dephosphorylation* of the C-terminal serine residues of TFEB in a similar manner as has been shown with other mTORC1-regulated N-terminal serine sites (S122, S142, S211) in cells. Moreover, we demonstrate that torin1-stimulated transcriptional activity of TFEB is highly preserved in AMPK null cells.

## Results and discussion

### AMPK is dispensable for mTOR-regulated (torin1-sensitive) transcriptional activity of TFEB

Previous studies, including our own, utilized a widely-used AMP-mimetic pro-drug AICAR to pharmacologically activate AMPK in cells, and reported that AICAR robustly promoted dephosphorylation and nuclear localization of TFEB [6,7]. However, AICAR has been shown to produce numerous off-target effects through regulating for example AMP/ZMP-sensitive enzymes [11–13]. We recently reported that AICAR treatment positively or negatively regulated (fold-change value of >1.3) a large proportion of differentially expressed genes [1026 out of 2053 in mouse embryonic fibroblasts (MEF) and 754 out of 1718 in mouse primary hepatocytes, respectively] in an AMPK independent mechanism [6]. In contrast, a benzimidazole derivative compound 991 (also known as ex229), a precedent structural analog of MK-8722, which potently activates all AMPK trimeric complexes through binding in a pocket termed allosteric drug and metabolite (ADaM) site (located at the interface of the AMPK α catalytic subunit and regulatory β subunit) [14,15] exhibited nearly exclusive specificity for targeting AMPK in both MEF and hepatocytes [6]. Consistent with our previous data [16,17], we verified that MK-8722 (1 µM) potently activated AMPK and did not affect any other kinases (>50% inhibition) in a panel of 140 human protein kinases in a cell-free assay (**Fig. S1A**).

We treated wild-type (WT) and AMPKα1/α2 double knock-out (DKO) MEF with MK-8722 (10 μ M) or torin1 (100 nM) for 2 or 6 h and assessed phosphorylation and transcriptional activity of TFEB by immunoblotting and reverse transcription quantitative PCR (RT-qPCR), respectively (**Fig. 1B and C, Fig. S1B and C**). Consistent with previous studies using 991[5,6], MK-8722 treatment caused a robust increase in phosphorylation of AMPK and its *bona fide* substrates acetyl-CoA carboxylase (ACC) and Raptor in WT, but not in AMPKα1/α2 DKO cells (**Fig. 1B, Fig. S1B**).

Torin1 treatment abrogated phosphorylation of an mTOR substrate p70S6K1 (S6K1) in WT and AMPKα1/α2 DKO, while MK-8722 elicited only a marginal inhibitory effect on S6K1 phosphorylation in WT (but not in AMPKα1/α2 DKO) cells (**Fig. 1B, Fig. S1B**). Both MK-8722 and torin1 treatment caused a profound dephosphorylation of TFEB at multiple established mTORC1-targeted residues (S122, S142, S211)[1] judged by phospho-specific antibodies and electrophoretic mobility band shift of total TFEB (and also TFE3) in WT. We next measured previously validated transcripts of torin1- or AMPK activator-sensitive TFEB target genes (*Hexa*, *Ctsa*, *Flcn*, *Fnip1*) [3,6,7] by RT-qPCR (**Fig. 1C and S1C**). There was no difference in the expression of the TFEB-targeted genes under basal/vehicle-treated condition between WT and AMPKα1/α2 DKO, except a modest decrease in *Flcn*. Both MK-8722 and torin1 significantly increased (∼1.5-2-fold) transcripts of all four genes 2 and 6 h following the treatment in WT cells (**Fig. 1C, Fig. S1C**). While MK-8722-stimulated increase in transcripts of TFEB target genes was abrogated in AMPKα1/α2 DKO, in sharp contrast to the previous study [7] torin1 induced comparable increases in the target gene expression between WT and AMPKα1/α2 DKO cells. To substantiate our findings using the cells genetically/chronically lacking AMPK, we also took a pharmacological approach and acutely inhibited AMPK by using a highly selective and potent inhibitor (BAY-3827) [18]. AMPKα1 and α2 were among the kinases most potently inhibited by BAY-3827, and a cell-free selectivity screen against 331 kinases revealed that BAY-3827 had better selectivity for AMPK [18] than previously described AMPK/ULK1 inhibitor SBI-0206965 [19,20]. However, it should be noted that ribosomal protein S6 kinases (RSK) 1-3 and their related kinases (MSK1, MST3) were identified as off-targets of BAY-3827 [18]. Thus, a proper negative control BAY-974, which is structurally related to BAY-3827 but has no inhibitory activity on AMPK [18], was also employed. We observed that BAY-3827, but not BAY-974, abrogated MK-8722-stimulated phosphorylation of ACC and Raptor, as well as dephosphorylation of S6K1 and TFEB/TFE3 **(Fig. 1D**). Both BAY-3827 and BAY-974 treatment showed no detectable changes in torin1-sensitive/mTORC1-regulated signaling or basal expression of the TFEB-target genes. BAY-3827 abrogated MK-8722-, but not torin1-stimulated induction of the TFEB-target gene expression (**Fig. 1E**). In contrast, but as expected BAY-974 treatment did not affect MK-8722- or torin1-stimulated gene expression. Taken together, loss of AMPK function via genetic or pharmacological means did not affect torin1-stimulated gene expression of the selected TFEB targets.

### Torin1-regulated TFEB/TFE3-dependent gene expression is profoundly preserved in cells lacking AMPK

TFEB plays a key role in maintaining cellular homeostasis through regulating expression of genes belonging to the CLEAR network [2,3]. To unbiasedly assess the requirement of AMPK in transcriptional activation of TFEB induced by pharmacological activation of AMPK or inhibition of mTORC1, we initially sought to establish MK-8722- and torin1-sensitive TFEB/TFE3-dependent genes through performing a transcriptome analysis using WT and TFEB/TFE3 DKO MEF (**Fig. 2A and C, Fig. S2A and B**). RNA-sequencing (RNA-Seq) analysis revealed that in WT MEF, 829 genes (15%) were commonly upregulated (FC ≥ 1.2, false discovery rate (FDR) < 0.05) in response to torin1 and MK-8722 compared to vehicle (DMSO) control and we observed a group of genes whose expression was downregulated in drug-treated TFEB/TFE3 DKO compared to WT MEF (**Fig. S2A and B**). To study this gene subset, differential expression analysis was performed and torin1 or MK-8722 stimulated genes that were also significantly downregulated (FC ≥ 1.2, FDR < 0.05) in TFEB/TFE3 DKO compared to WT MEF were subjected to enrichment analysis, and gene ontology functions were explored (**Fig. S2C and D**). This revealed that the downregulated genes in response to both torin1 and MK-8722 treatments in TFEB/TFE3 DKO compared to WT cells have cellular functions involved in autophagy, ion homeostasis, vacuolar transport and organization and lysosome organization and transport. As anticipated, some of the downregulated genes in torin1 *vs.* vehicle control in TFEB/TFE3 DKO compared to WT were found to be involved in regulatory functions of mTOR signaling (**Fig. S2C**). With MK-8722 treatment, endosomal transport functions were also enriched in downregulated TFEB/TFE3 DKO genes (**Fig. S2D**). Gene set enrichment analysis was performed on torin1- and MK-8722-treated cells compared to vehicle in TFEB/TFE3 DKO vs WT to study the top pathway associated with TFEB/TFE3 functions in these treatments, and this revealed that the highly enriched pathway for both MK-8722 and torin1 treatments is the lysosome (**Fig. S2E and F**). Furthermore, we observed that genes significantly downregulated (FC ≥ 1.2, FDR < 0.05) in TFEB/TFE3 DKO MEFs both torin1 and MK-8722 were commonly related to lysosomal and autophagy functions, confirming established key roles for TFEB/TFE3 in lysosomal functions regulated by mTORC1/AMPK signaling pathways.

**Figure 2.**
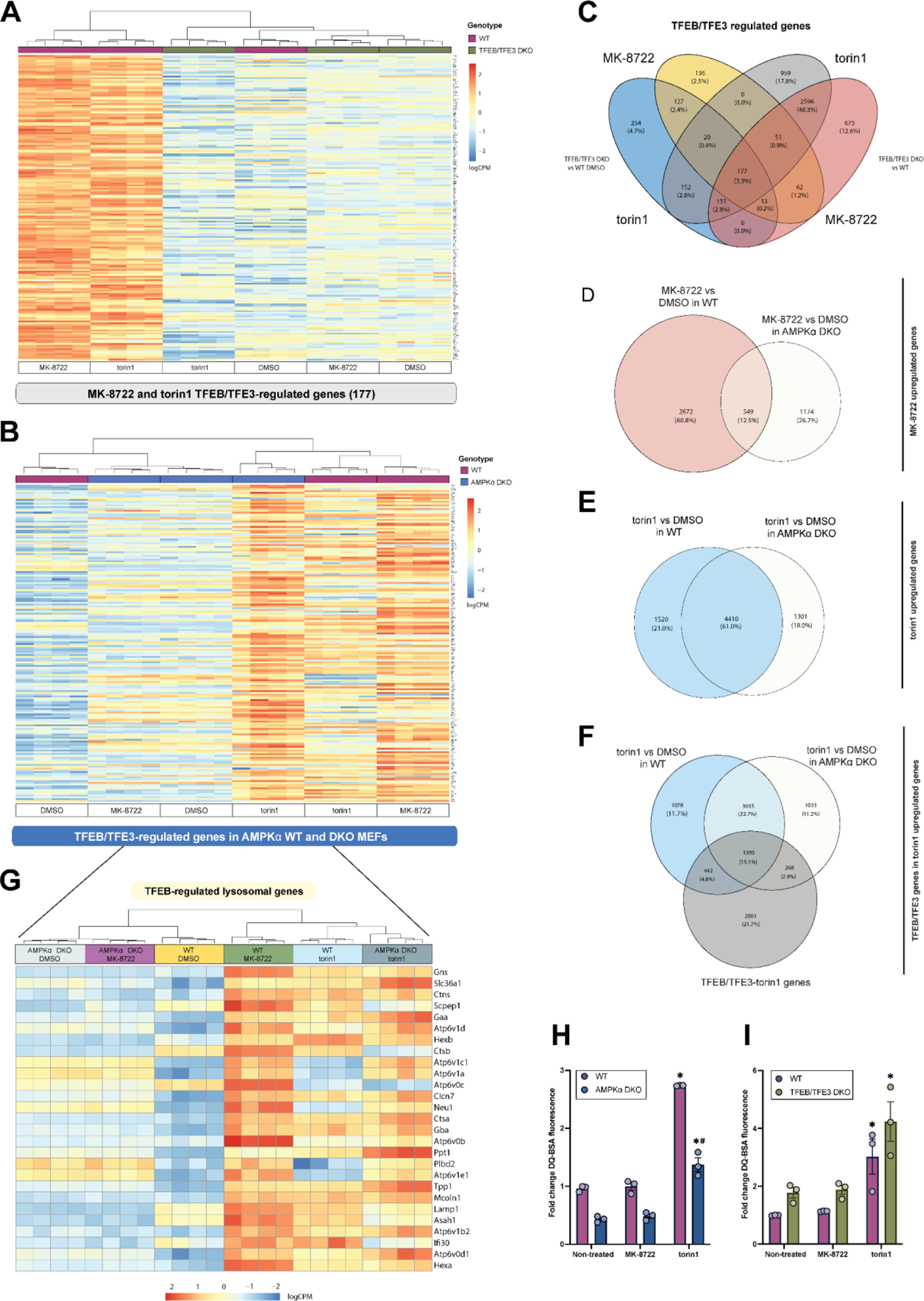
Unbiased messenger RNA sequencing substantiates dispensable role for AMPK in mTORC1-mediated/torin1-sensitive transcriptional activation of TFEB/TFE3. (A and B) Heatmaps of gene expression of MK-8722 and torin1-regulated genes in (A) WT and TFEB/TFE3 DKO MEF and (B) in WT and AMPKα1/α2 DKO MEF. (C-F) Venn diagrams depicting significantly altered transcripts across treatments, with (C) TFEB/TFE3-regulated genes by MK-8722 and torin1 were obtained by selecting significantly downregulated genes in TFEB/TFE3 DKO MEF cells compared to DMSO WT and across treatment conditions. Significantly upregulated genes upon MK-8722 (D) and torin1 (E) treatment across WT and AMPK DKO genotypes and (F) significantly upregulated TFEB/TFE3-genes present in torin1 upregulated genes. All significant genes were defined as having a fold change (FC) ≥ 1.2 and false discovery rate (FDR) < 0.05. (G) Heatmap of the previously reported TFEB-regulated lysosomal genes [3] present in current MK-8722- and torin1-regulated TFEB/TFE3-dependent genes. All expression data is shown as log2 counts per million (logCPM). (H, I) Lysosomal proteolytic activity expressed as fold change in DQ-BSA fluorescence in WT and (H) AMPKα1/α2 DKO MEF or (I) TFEB/TFE3 DKO MEF treated with vehicle (0.1% DMSO), 10 μM MK-8722 or 100 nM torin1 for 2 h. Representative data from three independent experiments are shown. Two-way ANOVA with Šídák’s multiple comparison was performed (* *p* < 0.05 vehicle *vs.* treatment and # *p* < 0.05 KO *vs.* WT).

To identify torin1- and MK-8722-sensitive TFEB/TFE3-dependent genes, we specifically focused on significantly downregulated genes in TFEB/TFE3 DKO compared to WT cells in these two treatments. The genes that fit this criteria and that were significantly downregulated in TFEB/ TFE3 DKO compared to WT MEF in both MK-8722 and torin1 treatments compared to DMSO vehicle were selected, and their expression profiles were visualized in a heatmap (**Fig. 2A and C**). This identified a total of 177 commonly downregulated genes in torin1- and MK-8722-treated TFEB/TFE3 DKO compared to the treated WT MEF (and from hereafter to be referred to as MK-8722- and torin1-sensitive TFEB/TFE3-dependent genes).

We then determined if the expression of TFEB/TFE3-dependent genes was altered in AMPKα1/α2 DKO MEF compared to WT in response to MK-8722 and torin1, and an unbiased heatmap was produced to visualize gene expression profiles (**Fig. S2G**). Differential expression analysis showed that 1007 (11.6%) and 807 genes (9.3%) were significantly upregulated (FC ≥ 1.2, FDR < 0.05) in response to torin1 and MK-8722, respectively, in WT MEF compared to vehicle (**Fig. S2H**). We then looked at the gene expression of TFEB/TFE3-regulated genes in AMPK WT/AMPKα1α2 DKO sequencing data using intact ENSEMBL IDs to select matching genes and visualized their expression profile in a heatmap (**Fig. 2B**). All TFEB/TFE3-regulated genes were matched in the AMPK WT/AMPKα1α2 DKO dataset except for the pseudogene Gm8615.

As anticipated, the majority (87.5%) of MK-8722-stimulated genes in WT were not upregulated in AMPKα1/α2 DKO cells, with only a 12.5% observed overlap (**Fig. 2D**). In contrast, a substantial 61.0 % of upregulated genes in response to torin1 treatment were conserved in both WT and AMPKα1/α2 DKO (**Fig. 2E**). Among those commonly upregulated genes, we observed that 15.1% (1395 genes) were predicted torin1-sensitive TFEB/TFE3-dependent genes **(Fig. 2G)**, including 27 genes (out of 46) that had been previously reported as direct TFEB target genes [3] (**Fig. 2C**). The depicted gene expression profiles revealed that the majority of MK-8722-stimulated genes were highly expressed in WT, but not in AMPKα1/α2 DKO cells, whereas torin1-stimulated genes were highly to moderately expressed in both genotypes. Collectively, unbiased transcriptome analysis revealed that AMPK is not essential for induction of vast majority of known TFEB/TFE3-regulated genes in response to torin1.

### TFEB/TFE3 and AMPK are dispensable for torin1-stimulated lysosomal proteolytic activation

It has been reported that both AICAR and torin1 acutely (within 2 hours) increase lysosomal proteolytic activity in WT MEFs, but not in AMPKα1/α2 DKO MEF [7] as measured by microscopic image-based DQ-BSA fluorescence, a self-quenched albumin probe that fluoresces upon degradation. We performed quantitative cytometry-based DQ-BSA fluorescence analyses in WT, AMPKα1/α2 DKO and TFEB/TFE3 DKO MEF treated with vehicle, MK-8722 or torin1 for 3 h (**Fig. 2H and I**). TFEB dephosphorylation, AMPK activation and mTOR inhibition were confirmed by immunoblotting (**Fig. S3A and B**). Even though lysosomal proteolytic activity was reduced in AMPKα1/α2 DKO cells compared to WT in basal (vehicle-treated) state as reported [7], we observed relatively similar increase (2-fold) with torin1 in both genotypes (**Fig. 2H**). MK-8722 had no effect on lysosomal proteolytic activity in both WT and AMPKα1/α2 DKO cells. Notably, we observed a modestly higher basal activity in TFEB/TFE3 DKO compared to WT cells, and torin1, but not MK-8722, had comparable increase in lysosomal proteolytic activity between the genotypes (**Fig. 2I**). This indicates that mTOR-dependent, but TFEB/TFE3-independent mechanism is responsible for torin1-stimlated lysosomal proteolytic activity in MEFs. Collectively, AMPK is required for basal proteolytic activity, but both AMPK and TFEB are dispensable for an acute torin1-stimulated lysosomal proteolytic activity in MEF.

### A synthetic peptide encompassing the TFEB C-terminal serine residues is a poor substrate of AMPK

A previous study using a cell-free assay showed that recombinant AMPKα1β1γ1 complex phosphorylated purified TFEB protein and that AMPK-mediated phosphorylation of the purified GST-TFEB C-terminal fragment (415-476) was modestly or profoundly reduced in the fragment harboring single serine to alanine substitution (S467A or S469A) or triple substitutions (S466A/S467A/S469A), respectively [7]. Consequently, a polyclonal phospho-specific antibody against a triple-phosphorylated peptide (p-S466, p-S467, p-S469) was developed, and the study showed that AICAR treatment significantly increased immunoprecipitation of TFEB/TFE3 against the p-S466/S467/S469 antibody compared to vehicle control in WT, but not in AMPKα1/α2 DKO cells [7]. However, specificity of the p-S466/S467/S469 antibody was poorly documented, and whether the antibody had exclusively detected the triple-phosphorylated protein or cross-reacted with single/dual-phosphorylated or non-phosphorylated protein was not reported. Notably, basal phosphorylation of p-S466/S467/S469 was similar between WT and AMPKα1/α2 DKO cells [7], indicating that these sites can also be phosphorylated by other kinases independently of AMPK. We developed individual single (in human TFEB p-S466, p-S467, p-S469 and in mouse TFEB p-S465, p-S466, p-S468) and dual p-S466/S467 (p-S465/466 in mouse) TFEB antibodies, and initially performed dot-blot analyses to assess specificity and sensitivity of the respective antibodies (**Fig. 3A**). To avoid confusion regarding numbering of the TFEB sequence (serine residues) between humans and mice, we hereafter use numbering for human TFEB sequence. To increase the chance of obtaining site-specific single p-S466 and p-S467 antibodies, we designed the antigen peptides where we placed p-S466 and p-S467 either at N-terminal or C-terminal end of the antigen peptide, respectively (**Fig. 3A**), so that phosphorylation of either site does not interfere detection of the other phospho-site. p-S466 antibody detected single p-S466 and dually phosphorylated p-S466/S467 peptide, and p-S467 antibody specifically detected single p-S467 phospho-peptide, whereas the p-S466/S467 antibody recognized dually phosphorylated p-S466/S467 and single p-S467 peptide with similar efficacy but cross-reacted to a lesser extent with single p-S466 peptide. P-S469 antibody specifically detected p-S469 peptide, but was less sensitive compared to other phospho-antibodies tested (**Fig. 3A**).

**Figure 3.**
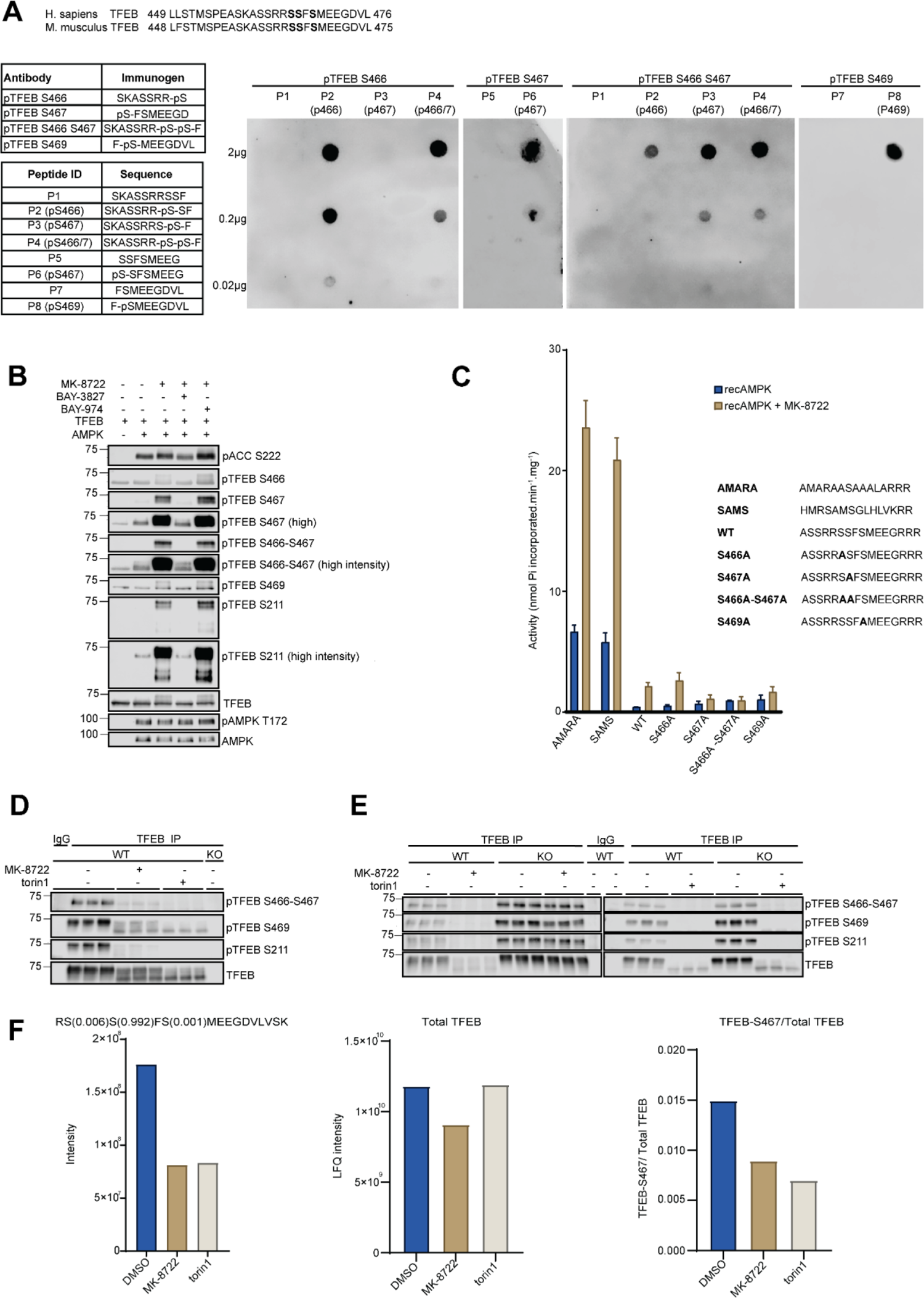
AMPK activation induces dephosphorylation of C-terminal phospho-sites of TFEB. (A) Sequence alignment of human and mouse TFEB. Sequence of immunogen used for generation of phospho-specific antibodies. Sequence of peptides used for dot blot analysis. Images of Dot blot assay for the phospho-TFEB-S466, -S467, -S466/S467 (dual) and -S469 antibodies. (B) Immunoblot images from *in vitro* phosphorylation of recombinant TFEB by recombinant AMPK in the presence of 1 μM MK-8722 co-incubated with 0.2 μM BAY-3827 or with 0.2 μM BAY-974. (C) *In vitro* AMPK activity assay in the presence of different peptides, including benchmarked peptide substrates SAMS and AMARA and peptides encompassing WT and mutated C-terminal serine residues of TFEB. (D and E) WT and AMPKα1/α2 DKO MEFs were treated with vehicle (0.1% DMSO), 10 μM MK-8722 or 100 nM torin1 for 1 hour. (D) TFEB/TFE3 DKO MEFs were used as controls. (D, E) TFEB was immunoprecipitated from protein lysates and C-terminal phospho-sites were detected by phospho-specific antibodies. pTFEB-S211 was used as a control. IgG was used as negative control for the TFEB immunoprecipitation. (F) TFEB-GFP knockin (KI) MEF were treated with vehicle (0.1% DMSO), 10 μM MK-8722 or 100 nM torin1 for 1 h. Pulldown TFEB-GFP by GFP-trap was analyzed by quantitative mass spectrometry. Phospho-peptide corresponds to TFEB (S467) quantitation based on intensity in the DMSO, MK-8722 and torin1 samples using data-dependent analysis. TFEB protein quantitation based on label-free quantitation (LFQ) in the DMSO, MK-8722 and torin1 samples using data-dependent analysis. TFEB phosphorylation in DMSO, MK-8722 and torin1 treated samples were quantified by taking ratio of the TFEB-S467 phosphorylation intensity with total TFEB quantitation.

Having established specificity of the antibodies, we evaluated phosphorylation of TFEB C-terminal serine residues by AMPK in a cell-free assay (**Fig. 3B**). As a control, we used bacterially expressed recombinant human ACC2 fragment (residues 29-249) and incubated it with recombinant AMPKα1β1γ1 complex in the absence or presence of MK-8722 (1 µM). This resulted in a robust increase in phosphorylation of ACC2, judged by signal intensity of a phospho-specific antibody against p-S222 or upward band mobility shift, which was particularly prominent in the presence of MK-8722. In contrast, AMPK only negligibly phosphorylated TFEB S466 even when its activity was enhanced (∼3-fold) by MK-8722 (**Fig. 3B**), whereas phosphorylation of S467 and S466/467 was modestly and robustly increased in the absence or presence of MK-8722, respectively. Notably, we also observed that phosphorylation of S211, an established mTORC1 phosphorylation site [21], was increased by AMPK to the similar degree compared to S467. In all the phospho-specific antibodies tested, MK-8722-stimulated phosphorylation was abrogated in the presence of BAY-3827 (0.1 µM), but not in the presence of the inactive compound BAY-974 **(Fig. 3B**).

In order to quantitatively assess phosphorylation of the TFEB C-terminal serine residues catalyzed by AMPK relative to benchmarked/optimized peptide substrates (SAMS, AMARA) [22,23] in a cell-free assay, synthetic peptides encompassing (surrounding) the C-terminal serine residues and its derivative peptides containing single or dual serine to alanine substitution, followed by an addition of three arginine residues to facilitate the binding of peptides to p81 phosphocellulose paper, were developed (**Fig. 3C**). In the absence of MK-8722, there was only negligible activity detected against TFEB C-terminal WT or S/A-substituted peptides. Even though MK-8722 enhanced activity against TFEB C-terminal peptides, it was ∼10-30 fold less compared to that against SAMS and AMARA peptides. These results are consistent with previous studies and bioinformatics motif prediction that all three of C-terminal serine residues of TFEB/TFE3 poorly match the AMPK substrate consensus motif [9,10].

### AMPK activation and mTORC1 inhibition promotes TFEB dephosphorylation at C-terminal serine residues

We assessed phosphorylation of C-terminal serine residues of TFEB in response to MK-8722 (10 µ M) and torin1 (100 nM) for 1 h in MEF. TFEB dephosphorylation, AMPK activation and mTOR inhibition in response to the treatment were confirmed in lysates by immunoblotting (**Fig. S3E and F**). We immunoprecipitated TFEB using a TFEB antibody (or IgG used as negative control) and blotted with p-TFEB antibodies. TFEB/TFE3 DKO MEF lysates were used as negative controls.

Consistent with our results shown earlier **(Fig. 1B**), both MK-8722 and torin1 caused robust dephosphorylation of TFEB S211 (**Fig. 3D and 3E**). In sharp contrast to the previous study [7], we observed that AMPK activation (induced by MK-8722), as well as mTOR inhibition (induced by torin1), resulted in a robust dephosphorylation of TFEB C-terminal serine sites. We confirmed that MK-8722-, but not torin1-mediated dephosphorylation of C-terminal serine residues of TFEB, is AMPK dependent (**Fig. 3E**) similar to the regulation of other known mTORC1 sites assessed earlier **(Fig. 1B**) or in previous studies [5,6]. To complement antibody-based approach, we performed mass spectrometry-based analysis of TFEB C-terminal serine phosphorylation. We generated C-terminally GFP-tagged TFEB knock-in (TFEB-GFP KI) MEF and treated them with vehicle, MK-8722, or torin1 for 1 h and immunoprecipitated TFEB-GFP protein using GFP-trap beads. The IPs were subjected to quantitative mass spectrometry analysis (**Fig. 3F and S3D, G and H**). We found that S467 relative to total TFEB quantitation was consistently dephosphorylated by ∼50% in response to both MK-8722 and torin1 and the data were confirmed by immunoblotting the IPs of TFEB-GFP by GFP-trap pulldown (**Fig. S3D**). Taken together, pharmacological activation of AMPK or inhibition of mTORC1 in cells commonly results in a robust dephosphorylation of TFEB not only at known mTOR-targeted N-terminal serine sites (S122, S142, S211), but also C-terminal serine sites. It remains unknown whether phosphorylation of C-terminal serine sites in TFEB is directly or indirectly regulated by mTORC1 [24] and also whether AMPK-dependent dephosphorylation of the C-terminal serine is mediated via dissociation of mTORC1 and TFEB from the lysosome through suppression of FLCN-FNIP1 GAP activity as proposed recently [5].

### Effect of alanine mutation of serine residue(s) in the C-terminal serine-rich motif on phosphorylation of other neighboring serine site(s)

Finally, we wanted to explore if loss or reduced phosphorylation of a specific site (or sites) in the C-terminal serine-rich motif of TFEB has impact on phosphorylation of other neighboring serine site(s). We ectopically expressed FLAG-TFEB WT and mutants (substitution of serine to alanine or threonine) in HEK293 cells and the lysates were subjected to immunoblot analysis with p-TFEB C-terminal serine antibodies (**Fig. S3C**). Consistent with the dot-blot analysis (**Fig. 3A**), we observed detection of p-S466/467 signal in WT but not in single S466A, S467A, double S466/467A or S466/467T mutant-expressing lysates. We also observed that S467A, but not S466A, showed a profound downward band shift. Notably, S469A, but not S469T, markedly increased p-S466/467 signal. Single S466A, S467A and double S466/467A reduced detection of p-S469 signal. Since both S466 and S467 residues are not included in the immunogen peptide (**Fig. 3A**), alanine mutation should not have interfered recognition by the p-S469 antibody. These results further validated specificity of the p-TFEB antibodies and implicated potential hierarchical phosphorylation. However, interpretation of results from mutagenesis experiments should be carefully assessed due to potential structural alternation intrinsic to artificial mutation (and C-terminal clustered serine residues of TFEB are within the disordered region [25]). It has been shown that introduction of triple S466/467/469A TFEB mutant blocks mTORC1-regulated TFEB activity [7], however it is unknown whether such effect was due to loss of phosphorylation or artefact caused by alanine mutations.

In conclusion, we demonstrate that AMPK activation and mTORC1 inhibition leads to dephosphorylation of conserved C-terminal serine sites in TFEB, and that AMPK is dispensable for torin1-sensitive transcriptional TFEB activation.

## Materials and methods

### Cell lines, culture and treatment

WT and AMPKα1/α2 DKO MEF [26,27] were provided by Benoit Viollet (Institut Cochin INSERM U1016). WT and TFEB/TFE3 DKO MEF [28] were provided by Rosa Puertollano (National Institutes of Health). HEK293 cells were purchased from Invitrogen (R75007). TFEB-GFP KI mice are generated by Taconic Biosciences through a constitutive KI of EGFP in the murine *Tfeb* gene via Easi-CRISPR/Cas9-mediated gene editing. The sequence for the open reading frame of EGFP was inserted between the last amino acid and the translation termination codon in exon 8. Spontaneously immortalized MEF were generated by Core Facility for Transgenic Mice (Department of Experimental Medicine, University of Copenhagen) according to the previously published protocol [29]. Cells were cultured in Dulbecco’s Modified Eagle Medium (DMEM [Invitrogen, 31966]) supplemented with 10% (v/v) fetal bovine serum (Sigma, F7524) and 1% (v/v) penicillin/streptomycin (Substrate and Sterile Centre, University of Copenhagen) in a 37°C incubator with 5% CO_2_. All cells were regularly tested and confirmed negative for the presence of mycoplasma throughout the experimental period. The following reagents were used for cell treatment: DMSO (Sigma, D4540), MK-8722 (Glixx Laboratories Inc, GLXC-11445 and MedChemExpress, HY-111363), torin1 (Selleck Chemicals, S2827), BAY-3827 (MedChemExpress, HY-112083) and BAY-974 (provided by Structural Genomics Consortium Toronto, ON, Canada).

### Transient transfection

HEK293 cells at 80% confluence were transiently transfected with the plasmids encoding FLAG-tagged TFEB or TFE3 using TransIT-X2 Dynamic Delivery System (Mirus, MIR6000). The next day after plating the cells in 6-well plates, the cells in one well were incubated with TransIT-X2:DNA complexes containing 3.5 µl TransIT-X2 reagent and 900 ng plasmid DNA in 250 µl Opti-MEM medium (Gibco, 51985-026). The cells were left for 18 h and then medium was replaced with full growth medium (DMEM supplemented with 10% FBS and 1% penicillin/streptomycin) for another 24 h followed by cell lysis. Following the treatment with the indicated drugs or vehicle, the cells were washed with room temperature PBS (Substrate and Sterile Centre, University of Copenhagen) and lysed in lysis buffer (50 mM Tris-HCl [Sigma, T1503], pH 7.5, 1 mM ethylenediaminetetraacetic acid disodium salt dihydrate (EDTA [Sigma, E5134]), 1 mM ethylene glycol-bis(2-aminoethylether)-*N,N,N′,N′*-tetraacetic acid (EGTA [Sigma, E3889]), 270 mM sucrose [Sigma, S7903], 1% (w/v) Triton X-100 [Sigma, X100], 20 mM glycerol-2-phosphate disodium [Sigma, G9422], 50 mM sodium fluoride (NaF [Sigma, 201154]), 5 mM sodium pyrophosphate decahydrate [Sigma, 221368], 1 mM DL-Dithiothreitol (DTT [Sigma, 43815]) containing protease and phosphatase inhibitors (500 uM phenylmethylsulfonyl fluoride (PMSF [Sigma, P7626]), 1 mM benzamidine hydrochloride [Sigma, 199001], 1 μg/ml leupeptin [Sigma, L2884], 1 μg/ml pepstatin A [Sigma, P5318], 1 μM microcystin-LR [Enzo Life Sciences, ALX-350-012], 1 mM sodium orthovanadate [Sigma, S6508]). The lysates were clarified by centrifugation at 6,000 x *g* for 10 min at 4°C and total protein concentration was determined using Bradford reagent (ThermoFisher Scientific, 23200) and BSA as standard. The lysates were snap frozen in liquid nitrogen and stored at −80°C till further analysis. The following plasmids were obtained from MRC PPU Reagents and Services (University of Dundee): pCMV5 TFEB 3xFLAG (identifier: DU67838), pCMV5 TFEB S466A 3xFLAG (identifier: DU73282), pCMV5 TFEB S467A 3xFLAG (identifier: DU73283), pCMV5 TFEB S466A S467A 3xFLAG (identifier: DU7328), pCMV5 TFEB S469A 3xFLAG (identifier: DU73878), pCMV5 TFEB S466T S467T 3xFLAG (identifier: DU80259), pCMV5 TFEB S469T 3xFLAG (identifier: DU80257).

### Western blotting

Proteins in the lysates were denatured at 100°C for 5 min in Laemmli sample buffer (50 mM Tris, pH 6.8, 2% sodium dodecyl sulfate (SDS [Sigma, 74255]), 0.5 mM EDTA (Sigma, E5134), 10% glycerol [Sigma, G5516], 0.01% bromophenol blue [Sigma, 114391]) and stored at −20°C until analysis. The denatured proteins were separated by SDS-PAGE on Mini-PROTEAN Tetra Cell (Bio-Rad Laboratories, Denmark) on 7-8% handcast mini polyacrylamide gels (acrylamide [Bio-Rad, 1610146], *N*,*N*,*N*′,*N*′-Tetramethylethylenediamine (TEMED [Sigma, T9281]), ammonium persulfate [Sigma, 248614], sodium dodecyl sulfate (SDS [Sigma, 74255]). The separated proteins were transferred from the polyacrylamide gel to the nitrocellulose membrane (Sigma, GE10600002) by Mini Trans-Blot Cell system (Bio-Rad Laboratories, Denmark). Membranes with proteins were blocked with 3% (w/v) skim milk (Sigma, 70166) in Tris-buffered saline (TBS, 20 mM Tris, 136 mM NaCl) containing 0.1% (w/v) Tween 20 (Sigma, P7949) (TBST). After blocking, the membranes with proteins were incubated with primary antibodies diluted in 4% bovine serum albumin (BSA [Sigma, A7030] in TBST containing 0.02% sodium azide [Sigma, 71290], overnight at 4°C. After washing with TBST, horseradish peroxidase (HRP)-conjugated secondary antibodies diluted in 3% skim milk in TBST were added to the membrane and incubated for 45 min at room temperature. The membranes were washed with TBST. The protein-antibody complex on the membranes were visualized by Odyssey XF Imager (LI-COR Biotech, LLC, Germany) after incubation of the membrane with the enhanced chemiluminescence substrate (Millipore, WBKLS0500), with an integration time of 0.5 min for most of the blots. For some proteins, a higher integration time was required, such as 2 min or 5 min.

The primary antibodies used were: ACC (Cell Signaling Technology, 3676; 1:1,000), p-ACC S79 (Cell Signaling Technology, 3661; 1:1,000), Raptor (Cell Signaling Technology, 2280; 1:1,000), p-Raptor S792 (Cell Signaling Technology, 2083; 1:1,000), AMPKα (Cell Signaling Technology, 2532; 1:1,000), p-AMPKα T172 (Cell Signaling Technology, 2535; 1:1,000), p70S6K (Cell Signaling Technology, 2708; 1:1,000), p-p70S6K T389 (Cell Signaling Technology, 9234; 1:1,000), TFEB (Bethyl Laboratories, A303-673A; 1:1,000), p-TFEB S122 (Cell Signaling Technology, 86843; 1:1,000), p-TFEB S211 (Cell Signaling Technology, 37681; 1:1,000), p-TFEB S142 (Millipore, ABE1971; 1:1,000), TFE3 (Cell Signaling Technology, 14779 and 81744; 1:1,000), tubulin (Sigma, T6074; 1:5,000). Affinity-purified p-TFEB S466 (immunogen sequence: SKASSRR-*p*S, where *p* denoted phosphorylated residue), p-TFEB S467 (immunogen sequence: *p*S-FSMEEGD), p-TFEB S466 S467 (immunogen sequence SKASSRR-*p*S-*p*S-F), p-TFEB S469 (immunogen sequence: F-*p*S-MEEGDVL) and p-TFE3 S321 (immunogen sequence: KAITVSN-*p*S-CPAELPN) were generated by Yenzym Antibodies, LLC and used at 1 µg/ml dilution. The secondary antibodies used were: HRP-conjugated anti-rabbit (Jackson ImmunoResearch Europe Ltd, 111-035-144; 1:10,000), HRP-conjugated anti-rat (Jackson ImmunoResearch Europe Ltd, 112-035-143; 1:10,000), HRP-conjugated anti-mouse (Bio-Rad, 1706516; 1:10,000).

### Dot blot

The phospho- and non-phospho-peptides encompassing the clustered C-terminal serine residues of TFEB/TFE3 were synthesized by GL Biochem (Shanghai) Ltd and had the following sequences: P1 (SKASSRSSF), P2 (SKASSR-*p*S-SF, where *p* denoted phosphorylated residue), P3 (sequence SKASSRS-*p*S-F), P4 (SKASSR-*p*S-*p*S-F), P5 (SSFSMEEG), P6 (*p*S-SFSMEEG), P7 (FSMEEGDVL), P8 (F-*p*SMEEGDVL). The peptides were spotted on nitrocellulose membranes and the membrane was left to dry for 30 min. The membranes were further blocked with 3% skim milk in TBST and the rest of the steps followed the protocol mentioned above in Western blotting.

### Immunoprecipitation

0.5 mg protein lysates were incubated with 20 µl 50% slurry Protein G Sepharose (Cytiva, 17061802) coupled with the relevant antibodies on an VXR basic Vibrax orbital shaker (IKA, Denmark) for 16 h at 4°C. At the end of the incubation time, Sepharose beads containing the immunoprecipitated protein were washed three times with lysis buffer supplemented with 500 mM NaCl (Sigma, S9888). The proteins captured on the beads were eluted from the beads by adding Laemmli buffer for 10 min at 65°C with vibration (1000 rpm). The eluents were subjected to SDS-PAGE as described above. The antibodies used for immunoprecipitation were: TFEB (Proteintech, 13372-1-AP; 2 µg antibody for 0.5 mg lysate) and negative control IgG from the same species (Cell Signaling Technology, 2729; 2 µg antibody for 0.5 mg lysate). For detection of protein-antibody complex, a light-chain specific secondary antibody (Jackson ImmunoResearch Europe Ltd, 211-032-171; 1:10,000) was used to avoid the potential interference with the heavy chain of the antibody.

### In vitro phosphorylation of TFEB by AMPK

Recombinant human AMPK (α1β1γ1 complex, SignalChem Biotech, P47-110GH) was diluted in enzyme dilution buffer (50 mM Tris, 0.1 mM EGTA, 1 mg/ml BSA, 1 mM DTT). 50 ng recombinant AMPK was pre-incubated with 0.2 µM BAY-3827 or 0.2 µM BAY-974 for 30 min and subsequently with 1 µM MK-8722 for 30 min. 250 ng recombinant TFEB (OriGene, TP760282) or recombinant ACC2 (Aviva, OPCD00449) was added to the reaction. The kinase reaction was performed in buffer A (50 mM HEPES [Sigma, H3375], pH 7.4, 0.1 mM EGTA, 100 mM MgCl_2_ [Sigma, M2670], 150 mM NaCl, 1 mM DTT) and initiated by addition of 100 μM ATP (Sigma, A1852) dissolved in 25 mM sodium borate (Sigma, 221732) and incubation at 30°C for 20 min with shaking (1000 rpm). The enzyme reaction was stopped by addition of Laemmli buffer and denatured at 100°C for 5 min. The samples were subjected to Western blotting analysis described above.

### In vitro AMPK activity assay

AMPK activity assay was performed as previously described [30]. Recombinant human AMPKα1β1γ1 trimeric complex (SignalChem Biotech, P47-110GH) was diluted in enzyme dilution buffer (50 mM Tris, 0.1 mM EGTA, 1 mg/ml BSA, 1 mM DTT) and pre-incubated with BAY-3827 (0.2 μM) or BAY-974 (0.2 μM) for 30 min and subsequently with MK-8722 (1 μM) for 30 min.

The kinase reaction was performed in buffer A (50 mM HEPES, pH 7.4, 0.1 mM EGTA, 100 mM MgCl_2_, 150 mM NaCl, 1 mM DTT) containing 100 μM ATP, 200 μM peptide and [γ-^33^P]ATP (Hartmann Analytic, SCF-301-12) at 30°C for 10 min. The enzyme reaction was stopped by spotting 75% of the reaction mixture onto cation-exchange P81 phosphocellulose paper (obtained from SVI Phosphocellulose (https://www.svi.edu.au/resources/phosphocellulose_paper/)) and the P81 phosphocellulose paper was washed in 75 mM phosphoric acid (Sigma, W290017), and then in acetone (Sigma, 179124). After washing, the radioactivity of the ^33^P incorporated into the substrate peptides was detected by Cerenkov counting in a Hidex 600SLe scintillation counter (Hidex Oy, Finland) for 5 min. The substrate peptides used were: AMARA (AMARAASAAALARRR, MRC PPU Reagents and Services) and SAMS (HMRSAMSGLHLVKRR, MRC PPU Reagents and Services). Peptides based on the C-terminal sequence of TFEB, TFEB C-terminal WT (ASSRRSSFSMEEGRRR), S466A (ASSRRASFSMEEGRRR), S467A (ASSRRSAFSMEEGRRR), S466A-S467A (ASSRRAAFSMEEGRRR) and S469A (ASSRRSSFAMEEGRRR) were obtained from GL Biochem (Shanghai) Ltd.

### RNA sequencing and bioinformatic analysis

Total RNA was isolated using QIAwave RNeasy kit (QIAGEN, 74536) with on-column DNAse treatment using Qiagen RNase-Free DNase set (QIAGEN, 79256) following manufacturer’s protocol. Messenger RNA sequencing was performed by the Single-Cell Omics platform at the Novo Nordisk Foundation Center for Basic Metabolic Research. Libraries were prepared using the Universal Plus mRNA-seq protocol (Tecan Life Sciences, 0520) as recommended by the manufacturer. Libraries were quantified with Qubit fluorometer (ThermoFisher Scientific, Waltham, USA), quality checked using a TapeStation instrument (Agilent Technologies, Waldbronn, Germany) and subjected to 52-bp paired-end sequencing on a NovaSeq 6000 (Illumina, San Diego, USA).

Data was processed using nf-core/rnaseq v3.11.2 of the nf-core collection of workflows [31]. The pipeline was executed with Nextflow v23.04.1 [32] with the reference genome set as *Mus musculus* mm10 GRCm38 release 102. Workflow parameters included STAR aligner and unique molecular identifiers (UMI) were considered using “--with_umi --skip_umi_extract --umitools_umi_separator:” parameters. Differential expression analysis was carried out in RStudio version 4.2.0 using the package edgeR and a batch-effect removal was performed using the package limma. Venn diagrams were produced using the R package ggvenn, with significant upregulated or downregulated genes described as FC ≥1.2 and FDR < 0.05. Heatmap figures of log2 counts per million (logCPM) computed using edgeR were produced with the R package pheatmap. Gene ontology analysis was conducted using clusterProfiler showing biological process (BP) ontology and a p-adjust method set to “BH” (Benjamini & Hochberg), with significant genes described as FC <u>≥</u>1.2 and FDR < 0.05. Gene set enrichment analysis (GSEA) was performed using gseaplot R package [33–37].

### qPCR

At indicated time points, cells were washed once with PBS (Substrate and Sterile Centre, University of Copenhagen) and lysed with Buffer RLT (QIAGEN, 79216) supplemented with β-mercaptoethanol (Sigma, M6250; 1:100 dilution). RNA was isolated, and on-column DNase treated with RNAse-Free DNase (QIAGEN, 79256), using RNEasy mini kit (QIAGEN, 74106) or QiaWave (QIAGEN, 74536) according to the manufacturer’s instructions. RNA concentration and quality was analyzed by a NanoDrop One Spectrophotometer (ThermoFisher Scientific, Wilmington, USA). 2 μg RNA was reverse transcribed using Superscript III cDNA synthesis kit (Invitrogen, 18080), dNTPs (Invitrogen, 10297-018), RNaseOUT (Invitrogen, 10777019), random primers (Roche, 11034731001) and nuclease free water (Ambion, AM9937) in a 20 μl reaction as per the manufacturer’s instructions. cDNA was diluted 20x and 2 μl was used for qPCR reactions (10 μl total) using indicated primers (Table in Supplemental Material) and PrecisionPLUS qPCR Master Mix with SYBRgreen (Primer Design, Z_PPLUS-SY-20ml) on a LightCycler480 (Roche Diagnostics A/S, Denmark). For each experiment, cDNA was diluted to generate a standard curve to analyze primer efficiency. The delta delta Ct method was used to calculate gene expression and data was normalized to mean of *Hprt* and *Tbp*.

### DQ-BSA assay

The cells were seeded on 12-well plates and the following day were loaded with 10 μg/ml DQ Red BSA (Invitrogen, D12051) in complete medium for 2 h at 37°C. At the end of the loading period, medium with DQ Red BSA was removed, and the cells were washed once with PBS and then treated with MK-8722 or with torin1 for 2 h. Cells were washed with PBS once and detached with trypsin (Gibco, 25300-054). Trypsinized cells were spun down at 500 x *g* for 5 min and cell pellets were resuspended for 20 min in 2% (v/v) formaldehyde (Sigma, 252549) solution in PBS for fixation at room temperature. Subsequently, cells were washed with PBS, spun down at 600 x *g* for 5 min and the resultant cell pellets were resuspended in PBS for analysis. Analytical flow cytometry was performed on CytoFLEX S cytometer (Beckman Coulter ApS, Denmark). Cell debris was gated out from total cells, and single cell gate was placed to analyse mean fluorescence intensity of the DQ-BSA dye by excitation with the yellow/green laser (561 nm) and detection in the channel with a band pass filter 610/20 nm. Data analysis was performed using FlowJo software (Tree Star).

### Immunoprecipitation followed by mass spectrometry

TFEB-GFP KI MEF cells were first lysed in NP-40 lysis buffer. Clarified lysates (60 mg protein) were incubated with GFP-Trap beads (30 μl packed beads) for 4 h on a rotating wheel at 4°C. Following incubation, beads were washed 3x with standard lysis buffer. Bead-bound proteins were denatured and eluted in 2x LDS for 5 min at 95°C. Samples were then filtered through Spin-X columns to remove the beads from the eluate. The filtered eluate was loaded onto a 4-12% Bis-Tris gradient gel and proteins were separated by SDS-PAGE. Gels were stained with InstantBlue and subsequently de-stained in deionised water. A small portion of the eluate was retained for analysis and validation by Western blotting. To minimise potential protein contaminants, all steps from this point were performed under a laminar flow hood. Disposable scalpels were used to cut protein bands of interest from the InstantBlue stained gels into 1-2 cm cubes, which were subsequently transferred into LoBind 1.5 ml Eppendorf tubes. Gel pieces were washed once in HPLC grade water, and then shrank in anhydrous acetonitrile (ACN) for 5 min with gentle shaking. The ACN was aspirated, and gel pieces were re-swollen with 50 mM Tris-HCl pH 8.0 for 5 min with shaking. The shrinking-swelling process was repeated once more, and the proteins within the gel pieces were reduced with 5 mM DTT in 50 mM Tris-HCl pH 8.0 for 20 min at 65°C. Next, the proteins within the gel pieces were alkylated with 20 mM iodoacetamide (IAA) in 50 mM Tris-HCl pH 8.0 for 20 min at room temperature in the dark. Gel pieces were then shrunk again in ACN for 5 min, dried and re-swollen in 50 μl of 50 mM triethylammonium bicarbonate (TEAB) pH 8.0 containing 5 mg/ml trypsin. After removing excess trypsin, gel pieces were covered in 50 mM TEAB and samples incubated in a shaker overnight at 37°C for tryptic digestion. An equivalent volume of ACN was added to the digest for 15 min with shaking and the supernatant was collected into a fresh LoBind 1.5 ml Eppendorf tube. Gel pieces were then re-swollen with 0.1% (v/v) trifluoroacetic acid (TFA) for 5 min with shaking, and peptides were extracted twice with ACN for 5 min each with shaking. After each extraction, the supernatant was collected and combined with the previous supernatants. The supernatants were then dried by vacuum centrifugation using a SpeedVac.

### Mass Spectrometry analysis

The total run time was set to 70 min. The mass spectrometer was operated in a data-dependent acquisition mode. A survey full scan MS (from m/z 350 to 1200) was acquired in the Orbitrap at a resolution of 60,000 (at 200 m/z). The automatic gain control (AGC) target for MS1 was set as 1.2 × 106, and ion filling time was set as 50 msec. The most abundant ions with charge state ≥ 2 were isolated in a 3-sec cycle, fragmented by using high-energy collision dissociation (HCD) fragmentation with 30% normalized collision energy, and detected at a mass resolution of 15,000 at 200 m/z. The AGC target for MS/MS was set as standard and ion filling time was set at custom, whereas dynamic exclusion was set to 30 sec with a 10-ppm (parts per million) mass window. These DDA data were searched using MaxQuant software. Protein sequence from Uniprot was used as a FASTA sequence to search the data. Trypsin was set as proteases with a maximum of two missed cleavages were allowed. Oxidation of Met, deamidation of Asn/Gln and phosphorylation of Ser/Thr/Tyr were set as a variable modification and Carbamidomethylation of Cys was set as a fixed modification. The default instrument parameters for MS1 and MS2 tolerance were used, and the data was filtered for 1% PSM, peptide and Protein level FDR.

### Statistical analysis

All experiments were independently performed at least 2-3 times with duplicates or triplicates for each experimental condition. The results are showed as mean ± standard error of the mean (SEM). Statistical significance was evaluated by two-way ANOVA in GraphPad Prism 10. Multiple comparisons test was performed by Šídák’s test. Statistical significance is indicated in the figure legends.

## Acknowledgements

We acknowledge Lars Roed Ingerslev, Thomas Gade Koefoed and the Single-Cell Omics platform at the Novo Nordisk Foundation Center for Basic Metabolic Research for technical and computational expertise and support. We thank the Core Facility for Flow Cytometry and Single Cell Analysis (University of Copenhagen) and the MRC-PPU Mass Spectrometry Facility for help with phospho-site mapping and MRC PPU Reagents and Services for DNA cloning. This work is funded by the Novo Nordisk Foundation Center for Basic Metabolic Research (NNF CBMR) based at the University of Copenhagen. NNF CBMR is an independent Research Center and partially funded by an unconditional donation from the Novo Nordisk Foundation (Grant number NNF18CC0034900 and NNF23SA0084103). C.F.B. is the recipient of a fellowship funded by NNF as part of the Copenhagen Bioscience PhD Programme, supported by grant (NNF20SA0035586). GPS is supported by the UKRI Medical Research Council (grant MC_UU_00018/6) and the pharmaceutical companies supporting the Division of Signal Transduction Therapy (Boehringer Ingelheim, GlaxoSmithKline, Merck-Serono).

## Abbreviations

AMPK: 5’-adenosine monophosphate-activated protein kinase

ACC: acetyl-CoA carboxylase

AICAR: 5-aminoimidazole-4-carbox-amide ribonucleotide

CLEAR: coordinated lysosomal expression and regulation

DKO: double knockout

DMEM: Dulbecco’s modified Eagle’s medium

DMSO: dimethyl sulfoxide

DQ-BSA: self-quenched BODIPY® dye conjugates of bovine serum albumin

KI: knock-in

KO: knockout

MEF: mouse embryonic fibroblasts

mTORC1: mechanistic target of rapamycin protein complex 1

RagC: Ras-related GTP-binding protein C

Raptor: Regulatory-associated protein of mTOR

RSK: Ribosomal S6 Kinase

RT-qPCR: reverse transcription quantitative polymerase chain reaction

S6K1: ribosomal protein S6 kinase 1

TFE3: transcription factor binding to IGHM enhancer 3

TFEB: transcription factor EB

ULK1: unc-51 like autophagy activating kinase 1

WT: wild-type

## Author contributions

Conceptualization: K.S. Experimental design: F.N., C.F.B., K.H., K.M.L., J.F.Z., G.S., K.S. Experimental execution: F.N., K.H., K.M.L., H.L., J.C., J.F.Z., G.S. Data analysis: F.N., C.F.B., K.H., K.M.L., H.L., J.C., J.F.Z., G.S., G.S., K.S. Data visualization and figure formatting: F.N., C.F.B, H.L. Supervision: G.S., K.S. Writing-Original draft preparation: K.S. Writing-Reviewing and Editing: All authors.

