## Supplementary Material for "AMPK activation promotes transcriptional activation of TFEB through its dephosphorylation"

| Gene | Forward 5-3 | Reverse 5-3 |
| --- | --- | --- |
| *Tbp* | CCTTGTACCCTTCACCAATGAC | ACAGCCAAGATTCACGGTAGA |
| *Hprt* | GCCCCAAAATGGTTAAGGTT | CAAGGGCATATCCAACAACA |
| *Hexa* | GCTGAGGGCACGTTCTTTATC | GCGAGATGTATCCAGCAGTACG |
| *Ctsa* | GAGCAGAACGACAACTCCCT | TGCCCACAATTCGAGACACT |
| *Flcn* | TGGATCGGATCTACCTCATCA | TGGACATCCAAACTGCTCTG |
| *Fnip1* | GATGCGTGTTCATGTCAAGG | GGAGAGTGGGTGCTTGCTAC |

**Supplemental Table 1**. Primer sequences used for this study.

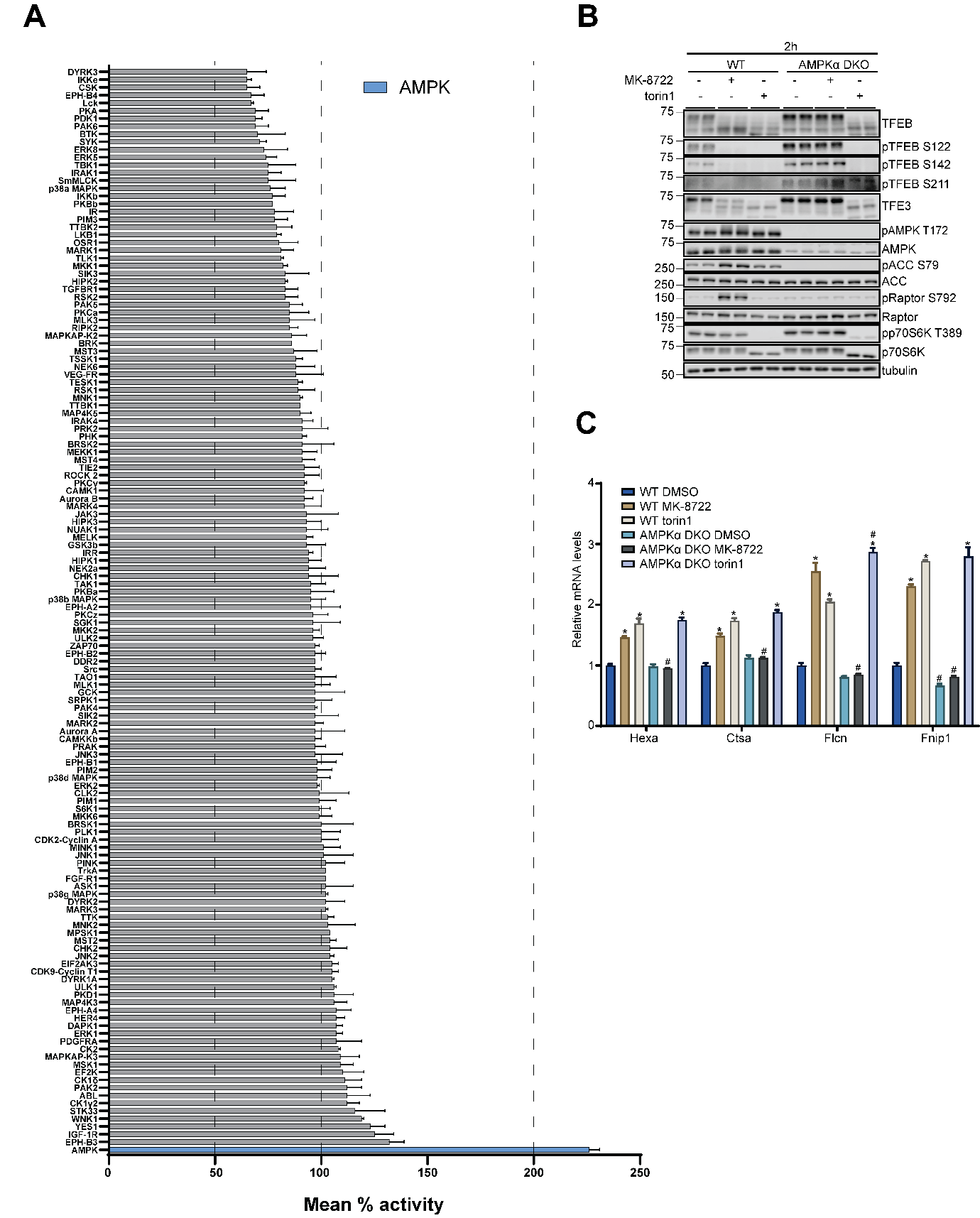

**Supplementary Figure S1**. Kinase selectivity profile for MK-8722 and AMPK-induced TFEB activation. (A) A screen of 140 human protein kinases (n = 2 per kinase, with or without 1 μM MK-8722) was performed *in vitro* using the MRC-PPU Premier Screen service as described in Methods. Results are expressed as mean ± SEM. AMPK (positive control) is shown in blue. (B and C) WT and AMPKα1/α2 DKO MEF were treated with vehicle (0.1% DMSO), 10 μM MK-8722 or 100 nM torin1 for 2 h followed by cell lysis for protein and RNA extraction. (B) Protein extracts were subjected to immunoblot analysis using the indicated antibodies. Representative immunoblot images from three independent experiments are shown. (C) The mRNA levels of the indicated genes were detected by RT-qPCR. Data from one independent experiment is shown. A two-way ANOVA with Šídák’s multiple comparison was performed (* *p* < 0.05 vehicle vs. treatment and # *p* < 0.05 KO vs. WT).

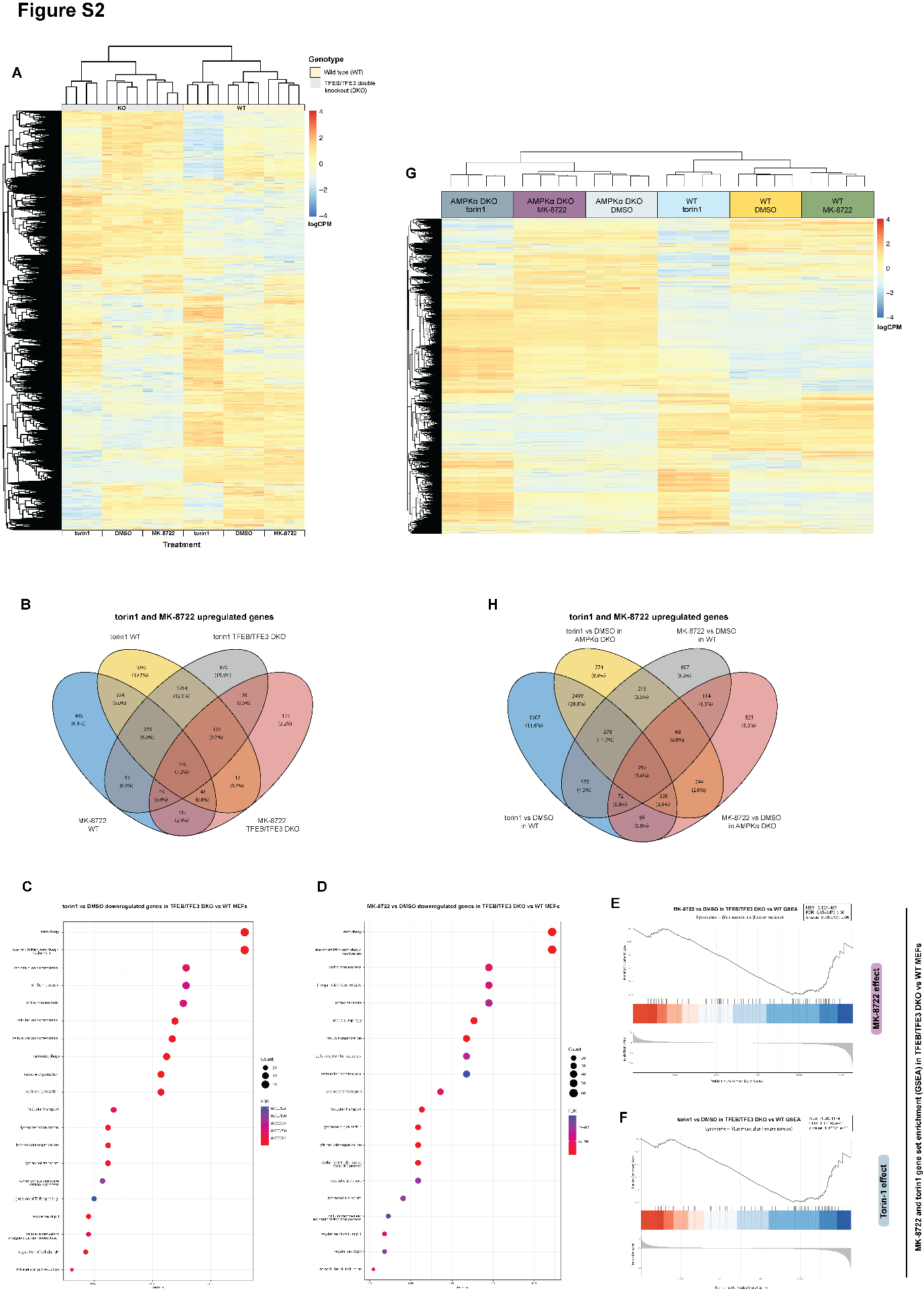

**Supplementary Figure S2**. Unbiased mRNA sequencing in wild type and double knockout (DKO) AMPKα or TFEB/TFE3 mouse embryonic fibroblasts treated with MK-8722 or torin1 reveals drug-regulated TFEB/TFE3 genes are associated with lysosomal functions. (A) Heatmap showing the gene expression profile of WT and TFEB/TFE3 DKO MEF treated with vehicle (0.1% DMSO), 100 nM torin1 or 10 μM MK-8722 shown in log2 counts per million (logCPM). (B) Venn diagram representing significant upregulated genes defined by FC ≥ 1.2 and FDR < 0.05 across torin1- and MK-8722-treated cells in both genotypes compared to vehicle. Gene ontology enrichment analysis was conducted on the significant downregulated genes in TFEB/TFE3 DKO compared to wild type in torin1 vs vehicle (C) and MK-8722 vs vehicle conditions (D), with the top 20 biological processes (BP) categories shown alongside gene count and FDR values. Gene set enrichment analysis (GSEA) was performed on vehicle vs MK-8722 (E) and vehicle vs torin-1 (F) in TFEB/TFE3 DKO vs WT conditions, looking at the TFEB/TFE3 DKO effect compared to WT, with the running enrichment score for the top pathway in both contrasts shown accordingly. (G) Heatmap showing gene expression profile in log2 counts per million (log2CPM) of WT and AMPK DKO MEF treated with DMSO, torin1 or MK-8722, with significantly upregulated (FC ≥ 1.2 and FDR < 0.05) gene numbers in torin1 and MK-8722 treatments compared to vehicle represented in a Venn diagram (H).

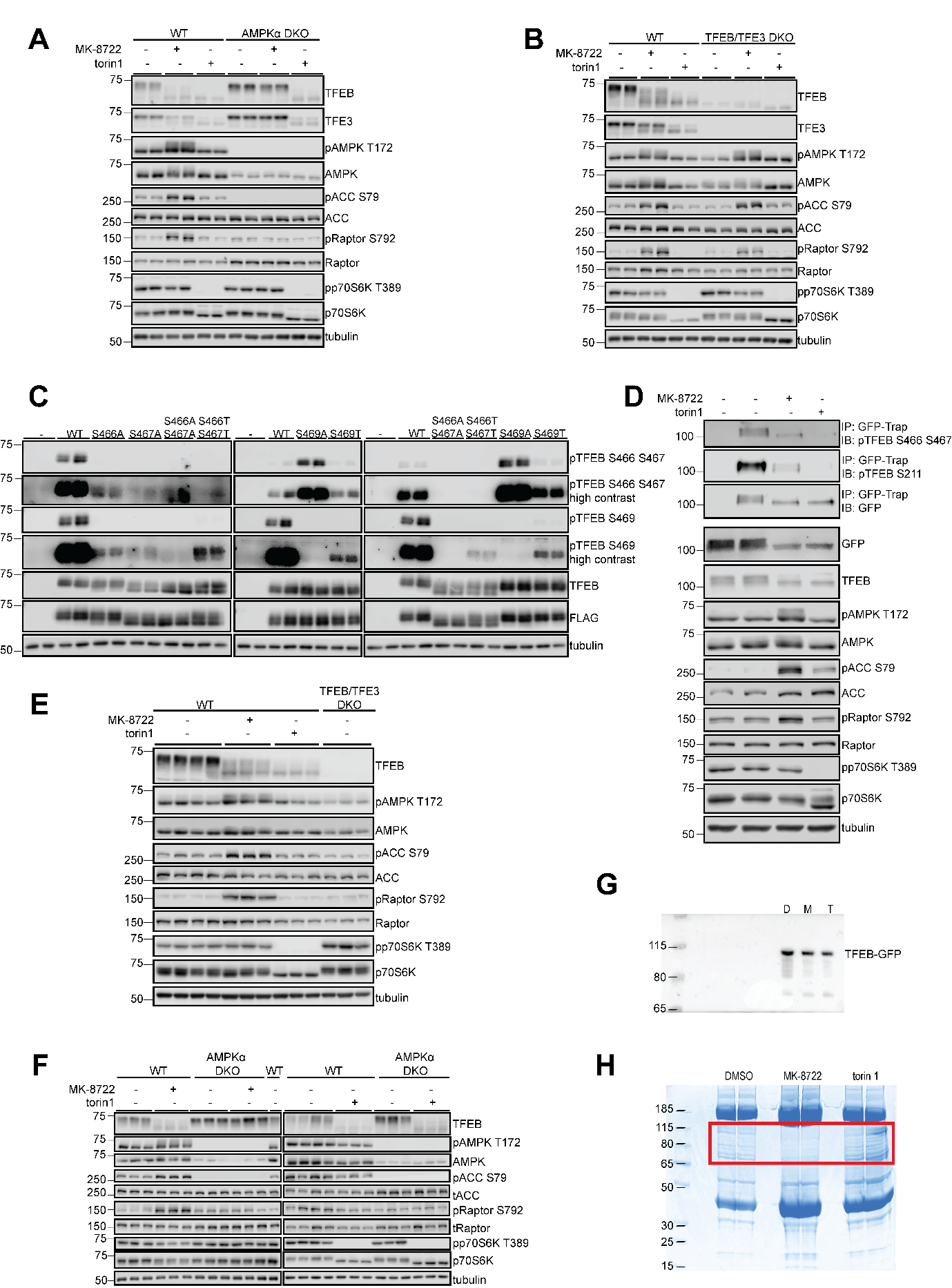

**Supplementary Figure S3**. AMPK activation induces dephosphorylation of C-terminal phospho-sites of TFEB. (A, B, E and F) WT, TFEB/ TFE3 DKO (A and E) or AMPKα1/α2 DKO (B and F) MEF were treated with vehicle (0.1% DMSO), 10 μM MK-8722 or 100 nM torin1 for 3 h (A and B) or 1 h (E and F) as controls for the experiments presented in Fig. 2H and 2I and in Fig. 3D and 3E. Protein lysates were collected at the end of the incubation time. The proteins indicated in the figures were detected by immunoblotting in isolated protein lysates. Representative immunoblot images from three independent experiments are shown. (C) HEK293 cells were transiently transfected with WT and different serine to alanine or threonine mutants and phosphorylation of TFEB was detected with different phospho-specific antibodies as indicated in the figure. (D) TFEB-GFP was immunoprecipitated by GFP-Trap from TFEB-GFP KI MEF treated with vehicle (0.1% DMSO), 10 μM MK-8722 or 100 nM torin1 for 1 h and the C-terminal phospho-sites depicted in the figure were detected with phospho-specific antibodies. (G and H) Detection of GFP by immunoblot and Coomassie blue gel staining in samples used for mass spectrometry analysis. The samples were pulldown TFEB-GFP from TFEB-GFP KI MEF treated with vehicle (0.1% DMSO) (D), 10 μM MK-8722 (M) or 100 nM torin1 (T) for 1h.
